# Cerebral blood volume sensitive layer-fMRI in the human auditory cortex at 7 Tesla: Challenges and capabilities

**DOI:** 10.1101/2022.08.02.502460

**Authors:** Lonike K. Faes, Federico De Martino, Laurentius (Renzo) Huber

## Abstract

The development of ultra high field (UHF) fMRI signal readout strategies and contrasts has led to the possibility of imaging the human brain in vivo and non-invasively at increasingly higher spatial resolutions of cortical layers and columns. One emergent layer-fMRI acquisition method with increasing popularity is the cerebral blood volume (CBV) sensitive sequence named vascular space occupancy (VASO). This approach has been shown to be mostly sensitive to locally-specific changes of laminar microvasculature, without unwanted biases of trans-laminar draining veins. Until now, however, VASO has not been applied in the technically challenging cortical area of the primary auditory cortex. Here, we developed a VASO imaging protocol for auditory neuroscientific applications. We describe the main challenges we encountered and the solutions we have adopted to mitigate them. With our optimized protocol, we investigate laminar responses to sounds. Finally, as proof of concept for future investigations, we map the topographic representation of frequency preference (tonotopy) in the auditory cortex.

**Highlights:**

- Layer fMRI VASO in the auditory cortex is challenging due to its physiology
- After protocol optimization we show the applicability of VASO to the auditory cortex
- Topographic maps obtained with VASO respect the large-scale tonotopic organization that has previously been shown with BOLD fMRI data.

## 1. Introduction

Ultra high field (UHF) magnetic resonance imaging allows the acquisition of functional data with increased sensitivity (Yacoub et al., 2001). This increased sensitivity can be used to breach into the mesoscopic scale in humans (for examples see De Martino et al., 2015; Huber et al., 2015; Kok et al., 2016; Lawrence et al., 2019; Nasr et al., 2016; Yacoub et al., 2007, 2008; Zimmermann et al., 2011), and the layered functional responses can be leveraged as a proxy for cortical architecture (Yang et al., 2021).

Gradient-echo blood oxygenation level dependent (GE-BOLD) functional magnetic resonance imaging (fMRI) is the conventional approach to collect submillimeter data, due to its relatively high signal-to-noise ratio (SNR) (Yacoub et al., 2005). However, T2*-weighted images collected at 7 Tesla (and higher fields) still contain contributions of both macro- and micro-extravasculature compartments (Uludağ et al., 2009; Yacoub et al., 2005). The macrovascular contribution to GE-BOLD originates from both pial vessels and draining vessels that penetrate the cortex orthogonally (Duvernoy et al., 1981). This results in two effects: the signal in superficial cortical depths is larger and the layer dependent spatial specificity is reduced as activation is drained away from the original locus of neural activity (Heinzle et al., 2016; Menon et al., 1995; Polimeni et al., 2016; Turner, 2002). Regardless, the increased sensitivity, coverage and temporal efficiency of GE-BOLD makes it the most common approach for laminar fMRI (for a recent review see e.g. De Martino et al., 2018), also when considering auditory studies (Ahveninen et al., 2016; Gau et al., 2020; Moerel et al., 2019; Wu et al., 2018).

While draining effects in GE-BOLD can be reduced with modeling and analyses approaches (see e.g. Havlicek et al., 2015; Markuerkiaga et al., 2016), alternative acquisitions have been proposed to minimize the contribution of macrovasculature. For example, spin-echo (SE) echo planar imaging (EPI) has been used to collect T2-weighted functional data (Yacoub et al., 2007, 2008). To retain T2-weighted specificity, these applications used segmented EPI acquisitions, while non segmented acquisitions introduce unwanted T2* contributions (Kemper et al., 2015). 3D gradient-echo and spin-echo (3D-GRASE - Feinberg et al., 2008; Oshio & Feinberg, 1991), has also been used to investigate human laminar and columnar function in both visual and auditory cortices (De Martino et al., 2013, 2015; Moerel et al., 2018; Olman et al., 2012; Zimmermann et al., 2011). However, the limited field of view (FOV) of early 3D-GRASE approaches has allowed only the investigation of small portions of cortex and, in auditory studies in particular, often in a single hemisphere (De Martino et al., 2015; Moerel et al., 2018; for a review see Moerel et al., 2021). More recent 3D-GRASE advancements can mitigate FOV constraints (Park et al., 2021). Furthermore a large spectrum of alternative approaches is currently under development to optimize the sensitivity and specificity of layer-fMRI experiments (Chai et al., 2020, 2021; Han et al., 2022; Kashyap et al., 2021; Kay et al., 2020; Lu et al., 2003; Priovoulos et al., 2022; Shao et al., 2021; Stanley et al., 2021; Truong & Song, 2009; Wang et al., 2021).

Cerebral blood volume (CBV) based imaging is one of the approaches to collect functional data with high spatial specificity, in which CBV functional responses can be acquired alongside conventional BOLD (Huber et al., 2019; Jin & Kim, 2008; Kim et al., 2013; Lu et al., 2013). The most commonly used approach to measure functional CBV changes is vascular space occupancy (VASO) (Hua et al., 2013; Huber et al., 2014; Lu et al., 2003). A concomitant acquisition approach of BOLD and VASO has the potential to combine their complementary aspects and facilitate a more comprehensive understanding of physiological underlying laminar activity changes. Furthermore, a combined acquisition of BOLD and VASO allows researchers to benefit from cumulative quality metrics of both methods, e.g. a high detection sensitivity (in BOLD compared to VASO) and a high localization specificity (in VASO compared to BOLD). VASO has been used to investigate laminar functional responses in visual (Huber et al., 2021a), motor (Huber et al., 2015, 2017), somatosensory (Yu et al., 2019) and prefrontal (Finn et al., 2019) cortices.

To date, VASO has not been successfully applied to investigate layer dependent functional responses in the human auditory cortex. Despite its lower power compared to BOLD (Beckett et al., 2020), the use of VASO has proven useful outside of auditory cortical areas (Finn et al., 2019; Huber et al., 2015, 2017; Huber et al., 2021a; Yu et al., 2019) and this warrants the need for developing an effective VASO protocol for auditory neuroimaging. Here, we present the results of a study aimed at optimizing a VASO functional protocol to image the auditory cortex at submillimeter resolution. We evaluated functional images collected at 7T using concurrent measurements of GE-BOLD and VASO. First, we explored a wide parameter space to mitigate methodological and physiological challenges. Specifically, we investigated the difference between a 2D- and a 3D-EPI readout and their stability across several participants. We showcase the laminar profiles of VASO data, and present initial results of the use of VASO for auditory neuroscience applications by characterizing VASO acquisitions of cortical sound frequency preference (i.e. tonotopic maps).

## 2. Methods

### 2.1 Ethics

The scanning procedures were approved by the Ethics Review Committee for Psychology and Neuroscience (ERCPN) at Maastricht University, following the principles expressed in the Declaration of Helsinki. Informed consent was obtained from all participants.

### 2.2 Participants

Participants were healthy volunteers with normal hearing and no history of hearing or neurological disorders. Participants were excluded if they had any standard MRI contraindications (e.g. any metal implants etc.).

Ten healthy volunteers participated in three separate studies. In study 1 (N=4), we optimized the VASO protocol. In study 2 (N=4), we evaluated the stability of the optimized protocol with 2D and 3D readouts. In study 3 (N=2), we applied the optimized protocol for tonotopic mapping as a proof of principle.

### 2.3 Scanner

Scanning was performed on a MAGNETOM “classic” 7T scanner (Siemens Healthineers) hosted by Scannexus (Maastricht) equipped with a 32-channel Nova Head Coil (Nova Medical, Wilmington, MA, USA). Sequences were implemented using the vendor provided IDEA environment (VB17A-UHF). We used an in-house developed 3^rd^ order B0-shim system (Scannexus) that depends on the vendor provided “3rdOrder ShimSet” feature.

### 2.4 Auditory stimulation

Sounds were presented to participants in the MRI scanner using MRI compatible ear buds of Sensimetrics Corporation (www.sens.com).

### 2.5 Slice-saturation slab-inversion VASO

We used a slice-saturation slab-inversion VASO (SS-SI-VASO - Huber et al., 2014) acquisition with either a 3D-EPI (Poser et al., 2010) or 2D-EPI readout (Huber et al., 2016). VASO uses an inversion recovery pulse to effectively null the contribution from the blood magnetization (Hua et al., 2013; Lu et al., 2003). For all of the tested protocols, the inversion delay (i.e. the dead time between the inversion pulse and the VASO signal readout module) was chosen to have the readout block roughly centered around the expected blood nulling time. In SS-SI-VASO, VASO and BOLD images are acquired in an interleaved fashion, which allows for a straightforward combination of the two datasets.

### 2.6 Reconstruction

The reconstruction of the data was conducted as described in previous studies for SMS-VASO (Huber et al., 2016) and 3D-EPI VASO (Huber et al., 2018b), respectively. In short, the vendor’s in-plane GRAPPA (Griswold et al., 2002) reconstruction algorithms were applied using a 3 × 2 (read direction x phase direction) kernel. Partial Fourier reconstruction (Jesmanowicz et al., 1998) was done with the projection onto convex sets (POCS) algorithm (Haacke et al., 1991) with 8 iterations. Finally, the complex coil images were combined using the vendor’s implementation of sum-of-squares.

SMS unaliasing was performed on-line on the scanner using a combination of the vendor software and the SMS reconstruction as distributed with the MGH blipped-CAIPI C2P (http://www.nmr.mgh.harvard.edu/software/c2p/sms). SMS signals were first un-aliased with an implementation of SplitSlice-GRAPPA with LeakBlock (Cauley et al., 2014) and a 3 × 3 SliceGRAPPA kernel before entering in-plane reconstruction as described above.

The 3D-EPI reconstruction was based on a previous 3D-EPI implementation (Poser et al., 2010) using a combination of standard scanner software and a vendor-provided work-in-progress implementation of GRAPPA CAIPIRINHA (Siemens software identifier: IcePAT WIP 571).

### 2.7 Study 1: Protocol optimization

We aimed at implementing and testing a VASO protocol for the auditory cortex that can mitigate a series of methodological challenges. The purpose of this pilot study was to explore the protocol parameter space of previously described 2D and 3D VASO sequences and optimize them with respect to maximal temporal signal-to-noise ratio (tSNR) and minimal artifact level. The protocol resulting from this study will then be subject to quantitative investigations and validations in a subsequent study (study 2).

First, compared to other cortical areas, the auditory cortex has an exceptionally short arterial arrival time of approximately 0.5-0.8s (see Fig. 7B in Mildner et al., 2014). This is approximately 1-2s earlier than the primary visual cortex. Such short arterial arrival times can result in the unwanted inflow of fresh (uninverted) blood during the VASO readout. When collecting simultaneous VASO and BOLD, these effects were more pronounced in the VASO data (figure 1A). To mitigate this challenge, we explored the usage of a phase-skipped adiabatic inversion pulse with B1-independent partial inversion (based on shapes of a TR-FOCI pulse - Hurley et al., 2010) that minimized these contaminants at the cost of SNR. Reducing the inversion efficiency by means of the phase skipped adiabatic inversion pulse can reduce the blood nulling time so that it is shorter than the arterial arrival time, mitigating inflow artifacts. Depending on the TR, the inversion efficiency and excitation flip angles that are used, the tissue signal can be reduced by about 30%.

**Figure 1.**
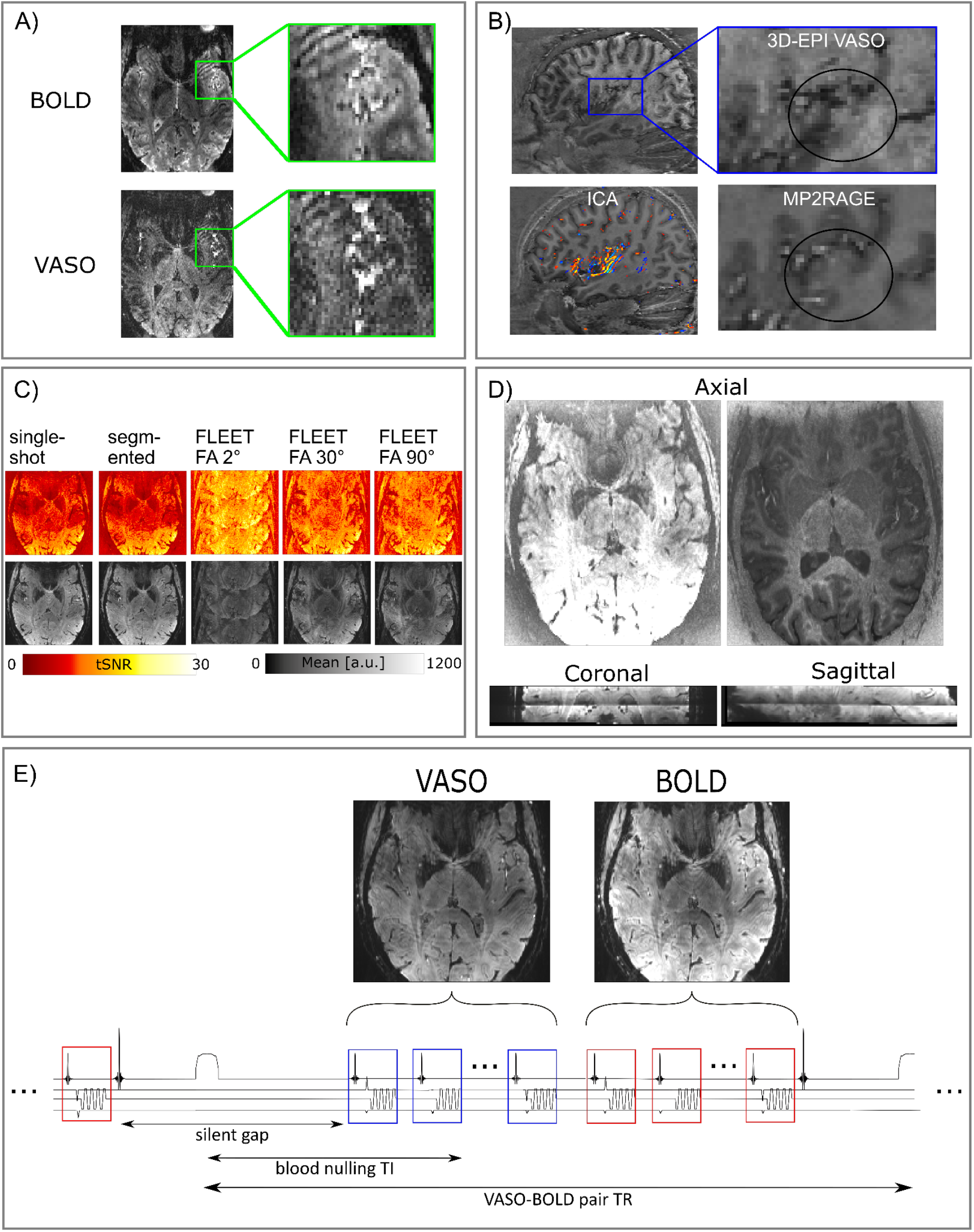
Overview of the challenges encountered when acquiring VASO data in the auditory cortex. A) Inflow effects were found in both GE-BOLD and VASO in temporal regions. However, the VASO signal seemed to be more affected by the inflow of not-nulled blood. B) Cardiac pulsation effects reduced image contrast due to long 3D-EPI readouts. In the functional images, the contrast in our region of interest seemed to be particularly affected. Additional ICA analysis (left bottom) showed the main components around Heschl’s gyrus. C) In the presence of physiological noise, there is a tradeoff in the amount of ghosts and the tSNR when evaluating different GRAPPA auto calibration signal (ACS) acquisitions. These tests were conducted for the protocol with 2D-SMS readouts. D) 2D-SMS VASO resulted in T1-weighted slice-wise intensity differences that were most visible in the middle of the slab. The two axial slices show the intensity differences between two “consecutive” slices (with the same signal intensity scaling). E) Schematic depiction of one TR of the final SS-SI-VASO sequence. An inaudible phase-skipped adiabatic pulse is used in the inherent silent gap of this sequence. This is followed by the acquisition of a volume of VASO and a volume of BOLD.

Second, we explored the effect that the readout time (and its relationship to the cardiac cycle) has on VASO data in the temporal cortex. Readout times longer than the cardiac cycle resulted in loss of contrast around Heschl’s gyrus (HG) and in typical vascular artifacts in components extracted with independent component analysis (ICA) from VASO time series (figure 1B).

High-resolution VASO is commonly used in combination with a 3D signal readout (e.g. 3D-EPI). However, since the primary auditory cortex, especially the medial portion of HG, is located right next to large feeding arteries, the partitioned 3D-EPI approach can result in higher susceptibility to physiological noise. To compare it to a 2D-EPI readout (study 2), optimizing parameters specific to the 2D readout was required. In particular, the location of the auditory cortex requires large in-plane imaging FOVs, resulting in large matrix size, and low bandwidth in the phase encoding direction for submillimeter acquisition protocols. The correspondingly long readout duration makes the acquisition protocol more susceptible to Generalized Autocalibrating Partially Parallel Acquisitions (GRAPPA, Griswold et al., 2002) artifacts. To find an effective protocol we compared the tSNR over 40 volumes resulting from an SS-SI-VASO acquisition with 2D readout at 0.9 mm isotropic employing different GRAPPA references: single-shot, segmented and FLEET (Polimeni et al., 2016) with three different flip angles (2, 30 and 90 degrees) (figure 1C). The segmented reference resulted in the best compromise between artifact level and tSNR in temporal areas.

Finally, we considered the use of 2D simultaneous multi slice (SMS - also known as multiband) (Moeller et al., 2010; Setsompop et al., 2012) EPI readouts in VASO in order to ‘freeze’ cardiac-induced vessel pulsation artifacts. The use of SMS results in different effective inversion times across slices and in our investigations this translated to sudden jumps of signal intensity in the VASO data (figure 1D). As this complicates the performance of retrospective motion correction and results in spatially heterogeneous tSNR we did not use SMS in the comparison in study 2.

Generally, in auditory fMRI studies, sounds are presented inside the silent gap between volume acquisitions (sparse design - Hall et al., 1999). In study 1, pure tones were presented for 800 ms within the inherent 900 ms dead time of the SS-SI-VASO sequence (thus following a sparse design), but this approach resulted in weak auditory evoked fMRI responses in the VASO (and simultaneously acquired BOLD) data. A possible reason for this reduced effectiveness of the sparse design is the relatively short duration of the gap and sound (900 ms and 800 ms respectively) compared to the noise of the BOLD/VASO acquisition time (∼2.5 seconds depending on the protocol). Following this rationale, in study 2 and 3 we continuously presented auditory stimuli (e.g. the auditory stimulation overlapped with the scanner noise) and played them loud enough to be audible compared to the scanner noise. This approach resulted in larger evoked responses (see results study 2 and 3).

A schematic depiction of the final protocol is illustrated in figure 1E (and with a complete parameter list available here: https://github.com/layerfMRI/Sequence_Github/tree/master/Auditory). In particular, we used an (inherently) inaudible adiabatic inversion pulse with a 30 degree phase skip, a readout time of 700 ms which is shorter than the cardiac cycle and a 70 degree reset-pulse (Lu, 2008) at the end of each acquisition of a VASO-BOLD pair. The purpose of the reset pulse was also to effectively saturate stationary Mz-magnetization of cerebrospinal fluid (CSF) and gray matter (GM) before the application of the consecutive inversion pulse. The suppressed CSF signal (see contrast in figure 1E) mitigates potential biases of dynamic CSF volume changes that have previously been reported to impose a source of bias for VASO applications in the auditory cortex (Scouten & Constable, 2007). The effective temporal resolution was 2.3 seconds.

### 2.8 Study 2: 2D versus 3D comparison

We collected two datasets of both BOLD and VASO (0.9 mm isotropic and 12 slices), one with a 2D readout (TR = 1833.5 ms; TE = 21 ms; flip angle = 70°; GRAPPA = 3; reference scan = segmented) and one with a 3D readout (TR = 1609 ms; TE = 22 ms; variable flip angles between 16° and 30°; GRAPPA = 3; reference scan = FLASH (Talagala et al., 2013)).

Participants were asked to passively listen to a series of sounds consisting of multi-frequency sweeps. Stimuli were presented following a blocked design with 20 volumes of sound stimulation followed by 20 volumes of rest. Each run consisted of thirteen stimulation blocks. A recording of the stimuli is available here: https://layerfmri.page.link/aud_stim. In each participant we collected two runs (S1 and S4) or 3 runs (S2 and S3) with a 2D readout and a 3D readout.

### 2.9 Study 3: Tonotopy

Simultaneous BOLD and VASO data were collected using the 3D sequence (after finalizing study 2) described above (0.9 mm isotropic; 12 slices; TR = 1609 ms; TE = 22 ms; GRAPPA = 3; reference scan = FLASH), variable flip angles between 16° (first segment of readout block) and 30° (last segment of readout block). In addition, we collected anatomical data (with optimized gray/white matter contrast) using MP2RAGE (TR = 6000 ms, TE = 2.39 ms, TI1/TI2 = 800/2750 ms, FA1/FA2 = 4°/5°, GRAPPA = 3 and 256 slices) (Marques et al., 2010) at a resolution of 0.7 mm isotropic.

Participants passively listened to tones varying slightly around 7 different center frequencies (130, 246.2, 466.3, 883.2, 1673, 3168 and 6000 Hz). Center frequencies were presented following a blocked design. Stimulation blocks (23 seconds) contained forty-six tones (500 ms each) varying 0.2 octaves around the center frequency. Each stimulation block was followed by a rest period (23 seconds). Functional runs consisted of fourteen stimulation blocks with a total duration of approximately 11 minutes per run. In one participant we collected four runs and in a second participant five runs. Before each tonotopic experiment tones were equalized for perceived loudness.

### 2.10 Functional data analysis

All functional images were sorted by contrast, resulting in a (BOLD-contaminated) VASO and a BOLD time series. The first three volumes of each time series were removed to account for the steady state. Each time series was motion corrected using SPM12 (Functional Imaging Laboratory, University College London, UK). The estimation of the motion parameters was restricted to a mask of the temporal lobe. Next, the time series were temporally upsampled by a factor of 2. This resulted in an interpolated TR of 1.15 seconds. As in previous studies, we corrected for the BOLD contamination in the VASO data using the open software suite LayNii (version 2.2.0 - Huber et al., 2021b).

In study 2, activation maps were created using AFNI (3dDeconvolve - version 21.2.04). We used a General Linear Model and normalized the time course with z-standardization. The resulting maps portray normalized differences between auditory stimulation and rest. Two-dimensional ROI’s were drawn manually in spatially upsampled EPI space and were divided in 7 equivolume layers (Waehnert et al., 2014) with which layer plots were created using LayNii.

In study 3, after preprocessing, functional data were first aligned to the anatomical data using Brainvoyager (version 22.2 - Brain Innovation, Maastricht, The Netherlands). For statistical analysis we used a General Linear Model with one predictor for each center frequency. Time series were normalized to percent signal change prior to statistical analysis. Tonotopic maps were created using best frequency mapping (Formisano et al., 2003).

## 3. Results

### 3.1 Study 2: 3D-EPI versus 2D-EPI

The presentation of auditory stimuli resulted in reliable responses in the bilateral auditory cortex for VASO (except for participant 2 in the 2D readout acquisition - see figure 2) and for BOLD (supplementary figure 1). For VASO, the 3D readout resulted in higher z-scores in bilateral auditory cortex, while this benefit was not directly visible in the BOLD data at these resolutions (supplementary figure 1). These results are somewhat consistent with previous 2D vs. 3D comparisons of VASO in the primary motor cortex (Huber et al., 2018a). Here we extend these findings for the physiological-noise constrained primary auditory cortex.

**Figure 2.**
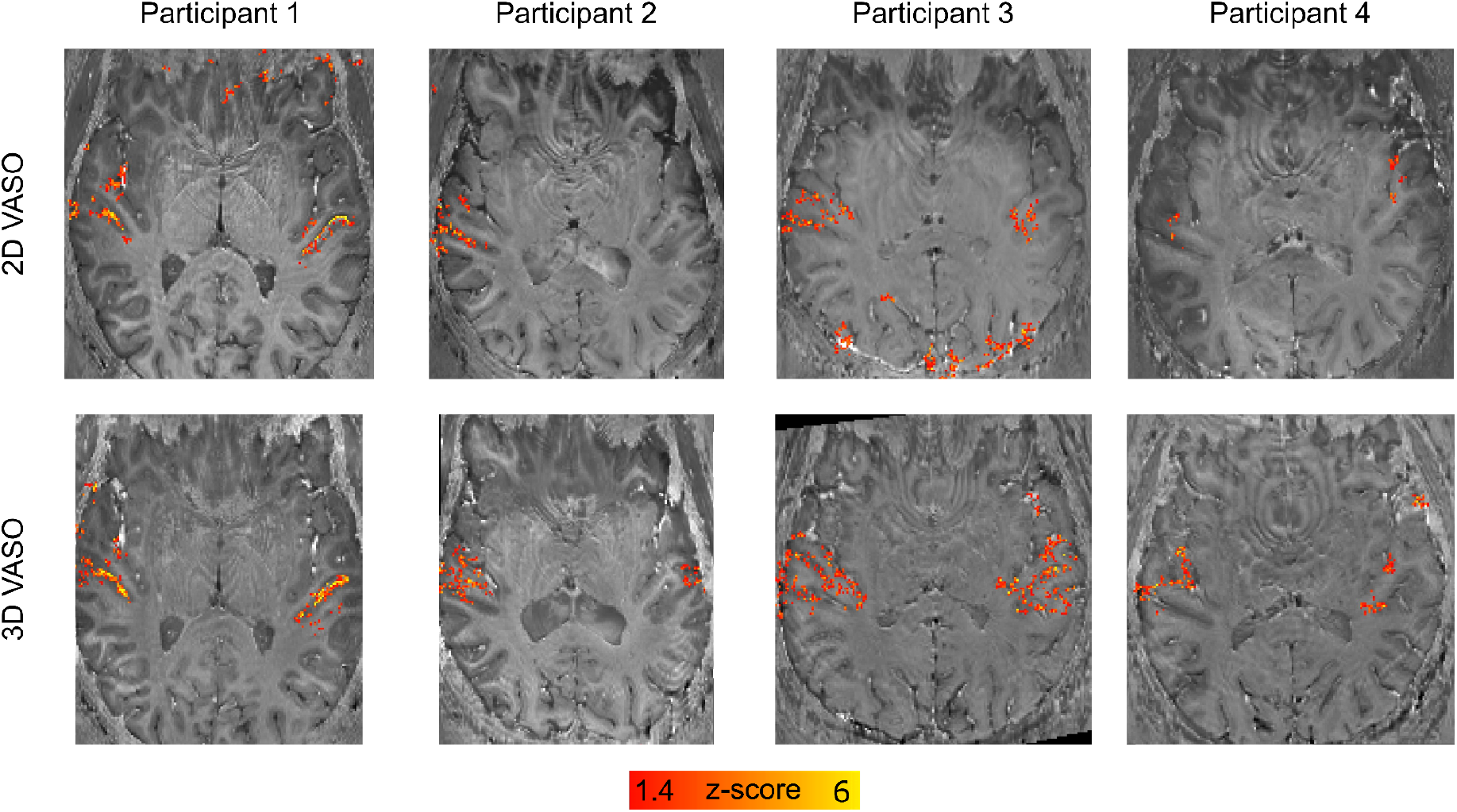
Activation maps of VASO (z-scores) overlayed on distortion corrected mean EPI images (per participant and readout). For our data, using a 3D-EPI readout seems to be beneficial in VASO.

Since the VASO signal is a composite signal from blood-nulled and not-nulled (BOLD) images, its detection sensitivity is indirectly dependent on the noise level of BOLD too. We believe, the result that VASO benefits from 3D-EPI more strongly than BOLD, is thus mostly driven by the relatively lower tSNR of blood-nulled images compared to non-nulled BOLD images.

### 3.2 Cortical depth-dependent responses

Figure 3 shows the layer profiles obtained in 2D regions of interest (ROIs; covering Heschl’s Gyrus [HG]). In VASO, the signal had a tendency to increase within gray matter. However, cortical depth dependent signal also showed a reduction at the pial surface (CSF/GM in figures 3 and 4), indicating its reduced sensitivity to macrovasculature. Separate analysis on the BOLD data using the same ROI definition, showed a monotonic increase in functional activation towards the cortical surface (supplementary figure 2) and no decrease on the pial surface. Similar results were obtained when defining ROIs based on functional activation (response to sounds) in the medial anterior part of HG on the left hemisphere (figure 4 for VASO and supplementary figure 3 for BOLD).

**Figure 3.**
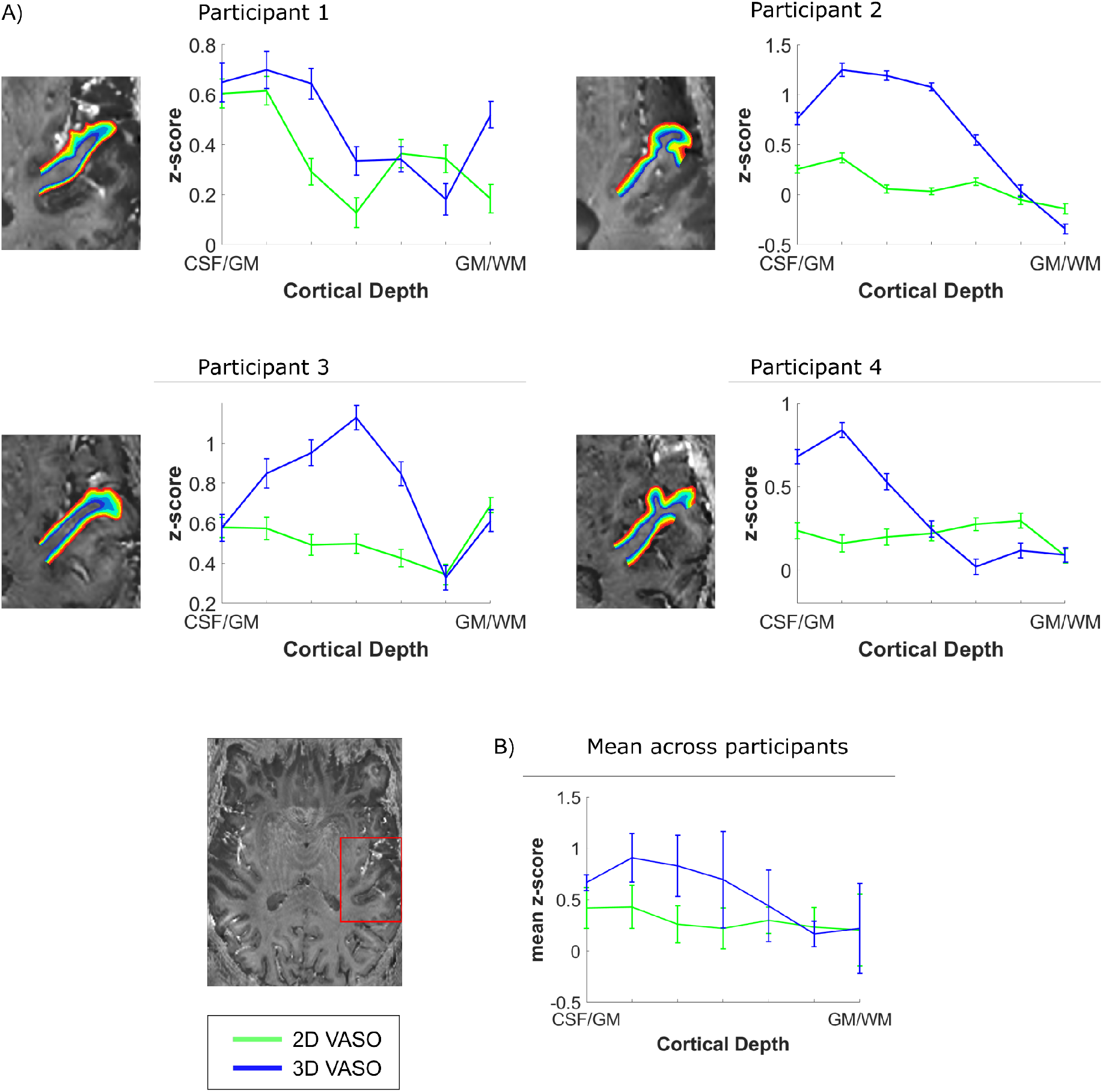
Z-scored cortical depth-dependent activation changes for the 2D- and 3D-EPI VASO data. A) The anatomically-informed ROI was drawn on the bias field corrected mean 3D-EPI VASO (example at the left bottom). The fMRI layer-dependent changes across depths for each participant. B) Average z-scored layer-dependent activation changes across participants.

**Figure 4.**
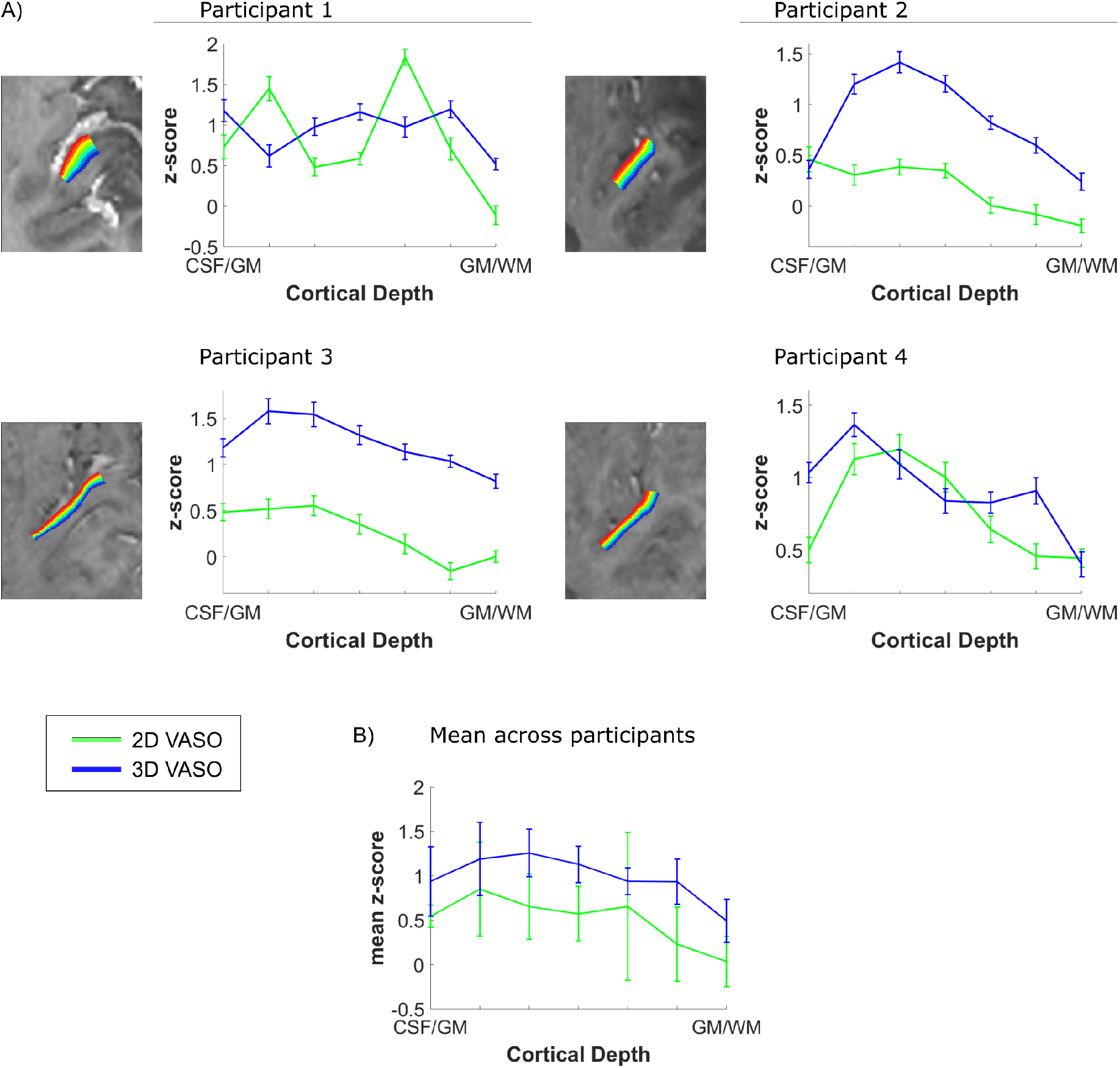
Z-scored cortical depth-dependent activation changes for the 2D-and 3D-EPI VASO data. The ROI was drawn on an axial slice as shown in figure 3A and was based on functional activation. A) The fMRI layer-dependent changes across depths for each participant. B) Average z-scored layer-dependent activation changes across participants.

Both the activation maps and the laminar analysis indicate that a 3D readout is beneficial for collecting VASO data (higher z-scores and increased reliability).

### 3.3 Study 3: Tonotopic maps

In study 3, the presentation of pure tones resulted in responses in the bilateral auditory cortex for both BOLD and VASO. Mid-gray matter anatomical surfaces were created from a WM/GM segmentation and inflated (figure 5) to visualize HG (outlined in black) and the planum temporale/polare. The analysis was confined to voxels showing both a positive signal change for BOLD (at a threshold of p<0.05 uncorrected) and a negative VASO signal change. Tonotopic maps that were interpolated on the mid-gray matter surface (figure 5) show the expected high-low-high frequency gradient along HG in VASO (see e.g. Moerel et al., 2014 for a comprehensive discussion on the expected topography of tonotopic maps). The same gradient is present in the BOLD data (supplementary figure 4) as shown in previous studies using GE-BOLD (Moerel et al., 2014).

**Figure 5.**
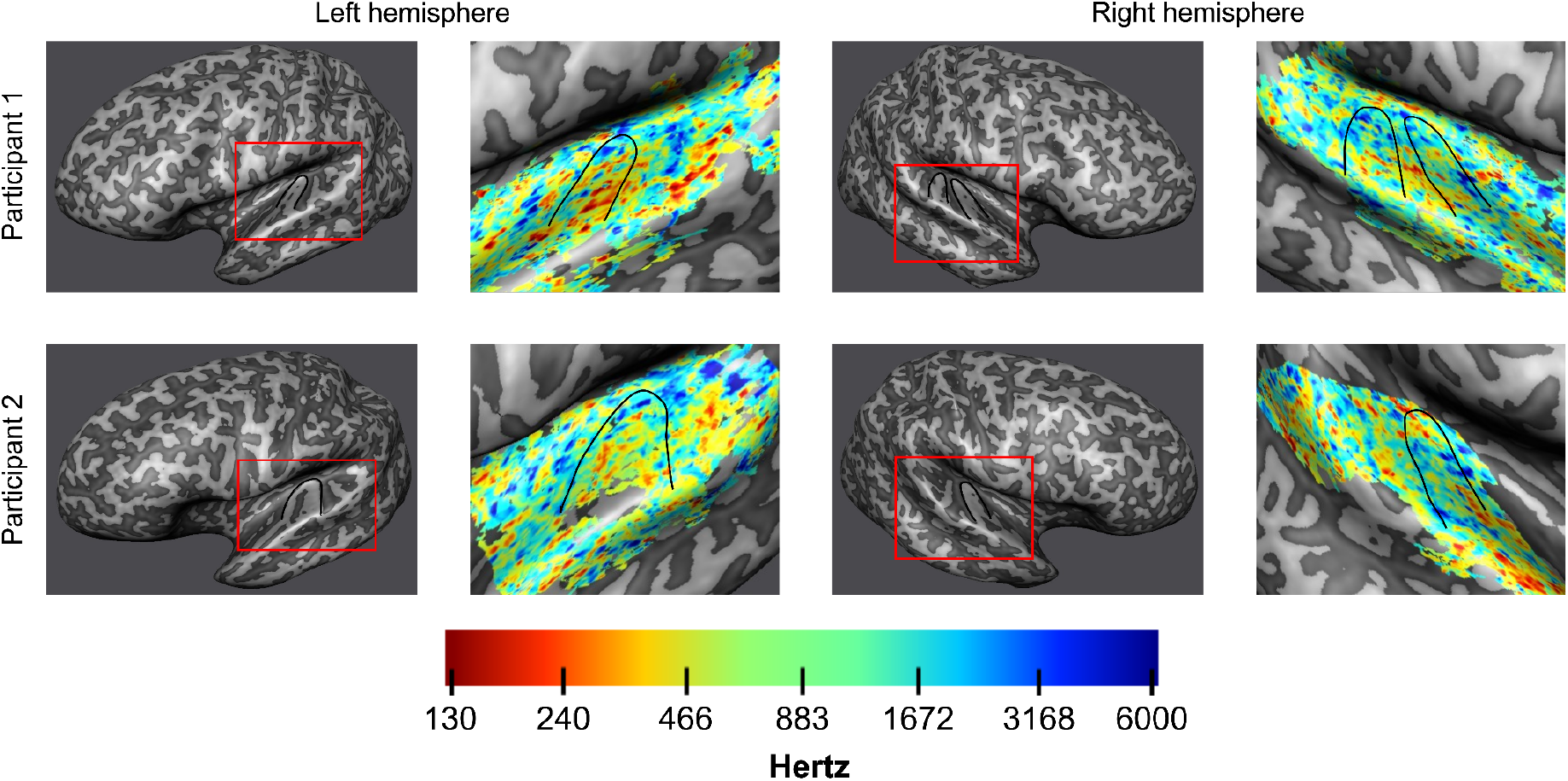
Inflated mid-gray matter surface meshes were created to visualize tonotopic maps coming from the VASO data. On the right of each inflated surface, tonotopic maps are displayed for both hemispheres of the two participants. Heschl’s gyrus is outlined in black. The expected tonotopic high-low-high frequency preference gradient is respected.

## 4. Discussion

Despite the fact that layer-fMRI VASO can provide valuable information in sub-millimeter and layer-fMRI applications (Finn et al., 2019; Huber et al., 2015, 2017; Huber, et al., 2021; Yu et al., 2019), it has not been successfully applied in the human auditory cortex. This is in contrast to GE-BOLD, which has been used to image laminar and columnar responses in the temporal lobe (Ahveninen et al., 2016; De Martino et al., 2015; Gau et al., 2020; Moerel et al., 2019). In this study, we aimed at implementing, testing, optimizing, and validating a VASO protocol for laminar fMRI investigations of the auditory cortices. As a proof of concept, we applied the final VASO protocol to map tonotopic responses in the bilateral auditory cortex.

Starting from a protocol that was previously successfully used (Huber et al., 2021b), the location and vascular physiology of the auditory cortex resulted in several artifacts. This required us to reconsider acquisition parameters and approaches that have helped to improve layer-fMRI applications but whose proof of generalizability across brain areas is still limited. The need to account for the specific vascular physiology of the auditory cortex, resulted in using a readout time shorter than the cardiac cycle and optimization of the inversion pulse (figure 1). To evaluate possible physiological contamination when using the standard 3D readout in VASO, we considered the use of a 2D readout. To develop an efficient 2D protocol for VASO, we combined techniques (such as FLEET for GRAPPA reconstruction and SMS acquisition) which are often used (in auditory neuroscience studies) when collecting submillimeter GE-BOLD data. To our surprise, while these approaches showed the expected utility in the GE-BOLD data, they did not result in the expected increase in sensitivity when considering the VASO data (figure 1). Study 1 resulted in two optimized protocols (2D and 3D readout) for VASO fMRI in the temporal lobe. Their comparison (study 2) showed an increased stability and SNR when using the 3D-EPI readout, despite its susceptibility to physiological noise. Note that the benefit for the 3D readout was particularly visible in the VASO time series (and not the BOLD data). It has been previously shown that the superiority of 2D-SMS or 3D-EPI readout strategies at 7T in conventional BOLD is highly dependent and specific to the acquisition and analysis details including the TR, acceleration factor, resolution, physiological noise correction and number of slices (Huber et al., 2018a; Jorge et al., 2013; Le Ster et al., 2019; Reynaud et al., 2017; Stirnberg et al., 2017). The BOLD results presented here are in agreement with this literature (see supplementary figures 1, 2 and 3).

We examined laminar profiles of activation elicited by the sounds presented in study 2. Similarly to previous studies investigating the specificity of laminar functional responses in auditory cortex (Moerel et al., 2018 - using 3D-GRASE), we did not observe a clear peak in functional response in middle cortical depths. The nature of the stimulation, and analysis could underlie this observation. First, as the auditory stimuli we presented in study 2 comprised complex dynamic sounds presented for about 20 seconds, it is unclear what the expected neural laminar profile would be in absence of any control for attention or another task. Second, we defined regions of interest for the laminar profiles based on macro-anatomy (anterior HG) or activation. The effect that this has on sampling the laminar activation profiles in auditory regions whose cytoarchitecture overlaps only partly with macro-anatomical features (see e.g. Gulban et al., 2020) is beyond the scope of this paper but could be an interesting venue for future investigations. What we did observe was that while the signal in the upper layers has the tendency to be larger than in middle and deeper layers, the signal decreases again at the pial surfaces. This is expected due to VASO’s insensitivity to large pial veins. As expected, GE-BOLD data resulted in an increased response towards superficial layers (supplementary figures 2 and 3) without a reduction on the pial surface. This profile is characteristic of GE-BOLD submillimeter acquisitions and is resulting from vascular draining and the contribution of large vessels on the cortical surface. If confirmed when analyzing a larger sample, a more controlled stimulus design, and within a more extended portion of temporal areas, the fact that vein-free VASO signal changes (Kim et al., 2013) within GM decrease as a function of cortical depth, could be interpreted as a validation of previous BOLD results (e.g. indicating that the signal trends visible in the BOLD signal in temporal areas cannot be solely explained by draining vein effects alone). It is important to note that while we here demonstrate that VASO auditory responses are not affected by draining and large vascular contributions on the cortical surface, we do not imply that in presence of careful controls (Kok et al., 2016; Lawrence et al., 2019; Moerel et al., 2018) or using modeling techniques (Havlicek et al., 2015; Markuerkiaga et al., 2016) GE-BOLD data cannot be used to investigate laminar cortical processing.

As a first proof of concept of the usability of VASO fMRI for the investigation of cortical processing in the temporal lobe, we presented results from a tonotopic experiment. Neurons throughout the auditory pathway display preferential tuning to the sound frequency (Winer & Schreiner, 2011) and using fMRI the topographic arrangement of frequency preference (tonotopy) can be mapped in single individuals (Ahveninen et al., 2016; Da Costa et al., 2015; Formisano et al., 2003; Moerel et al., 2014). Tonotopy shows a typical topography with a low frequency region residing primarily on the HG and regions preferring high frequency bordering it both posterior medially and anterior laterally (for a description see e.g. Moerel et al., 2014). This characteristic topography makes tonotopy a possible benchmark for auditory functional acquisitions. The large scale tonotopic gradient covering the superior temporal plane was visible in the VASO data. This initial promising result opens the venue to further investigations on the specificity of the VASO signal across cortical depths (Moerel et al., 2018).

In both study 2 and 3 we opted for a continuous stimulation protocol (i.e. with sounds overlapping with the scanner noise). This choice was driven by initial investigations (study 1) that showed weak effect size for a strategy more conventional in auditory fMRI (i.e. a sparse design - Hall et al., 1999). A possible reason for this is the relatively short duration of the silent gap inherent in the VASO acquisition (900 ms) compared to the scanner noise time (∼2.5 seconds), making a sparse design highly inefficient in our case despite its quiet stimulation characteristics. The acquisition protocol we provide here could serve as the basis of future investigations directed at optimizing sound presentation in auditory VASO fMRI. We nevertheless consider it reassuring that despite the fact that we opted for presenting sounds also simultaneously with the scanner noise, tonotopic maps resulting from the VASO (figure 5) and BOLD acquisition (supplementary figure 4) conform to the expected topography.

Despite the shown applicability of VASO for auditory fMRI, we deem it necessary to outline some limitations (many not specific to auditory studies) that require consideration when setting up a neuroscientific (laminar) fMRI study. While VASO is more sensitive to microvascular CBV increases, it is also characterized by a reduced detection sensitivity (as indicated by generally lower z-scores in figures 2, 3 and 4 than in supplementary figures 1, 2 and 3). To compensate for this effect a typical approach is to average across runs. As a result, extending averaging across sessions would require careful consideration of approaches for inter session alignment (and placement) of the relatively small slab (12 slices in our case). Future investigations may have to address issues related to detection sensitivity and its dependence on experimental design and sound presentation schemes. In addition, when using VASO, functional runs are typically acquired with an identical design as averaging is performed on the raw time series before BOLD correction to limit noise amplification. This calls for careful balancing of conditions within functional runs. To increase sensitivity we also employed long stimulation periods (block design). Evaluating the sensitivity of event-related functional responses with VASO (Dresbach et al., 2022) would increase its usability (e.g. to prediction-error related responses in typical oddball designs). Moreover, alternative approaches for increasing sensitivity such as denoising (e.g. NORDIC - Vizioli et al., 2021), should be considered in future investigations. Finally, while to compensate for physiological noise effects we decided to use a readout train shorter than the cardiac cycle, in the future it may be interesting to consider higher order physiological noise correction methods in k- space.

We believe that the significance of this work is multi-fold. In study 2 we developed a ready-to-use acquisition protocol for the user base of application-focused neuroscientists. The sequence binaries and the importable protocols are publicly available via ‘SIEMENS’ sequence ‘app-store’ on TEAMPLAY for any users of a ‘classical’ MAGNETOM 7T, which is the most widely used 7T scanner version around the world today. Users of other scanner versions and vendors will benefit from this protocol-development study as they can re-implement the acquisition approaches as described in study 1. Additionally, our protocols may allow application studies that are not straightforwardly addressable with the vein-bias of conventional GE-BOLD, such as single-task condition experiments. In such experiments, utilizing the developed VASO protocols alongside with BOLD can be useful to augment the understanding of the neurovascular origin of the fMRI signals. Other example studies, where acquiring VASO and GE-BOLD simultaneously may be beneficial, might be related to research questions of altered vascular baseline physiology (e.g. in studies about pharmacological interventions, aging, and surgical interventions). Furthermore, we think that the concomitantly acquired VASO and BOLD data can be useful to calibrate existing layer-fMRI BOLD models (Corbitt et al., 2018; Havlicek et al., 2015; Heinzle et al., 2016; Markuerkiaga et al., 2016; Marquardt et al., 2018; Merola & Weiskopf, 2018; Puckett et al., 2016) and extend their applicability across brain areas. For example, future GE-BOLD studies that want to apply venous-deconvolution model-inversion and may not find an increased response in the middle layers, can use the data we present here to increase the confidence in their results. The imaging protocol developed here can have implications beyond the auditory cortex. The auditory cortex is not the only brain area challenged by proximal macro-vessels with substantial physiological noise. There are many other brain areas in which sub-millimeter VASO was not successfully applied until now, for example, hippocampus, insular cortex, claustrum, entorhinal cortex, and thalamic nuclei. The protocol developed here is designed to mitigate the challenges of such brain areas and may be useful to address neuroscientific research questions in those brain areas as well. Finally, the main aim of this work was to provide the auditory research community with a viable VASO protocol for laminar fMRI studies.

To conclude, our results demonstrate that, when using optimized parameters, VASO can be used to investigate cortical responses in the bilateral temporal cortex. While VASO has a lower detection threshold compared to GE-BOLD, it is believed to be dominated by microvascular CBV increase close to the site of neural activity changes. A combined acquisition approach of BOLD and VASO, as described here, may allow benefitting from the quality features of each method.

## Acknowledgements

The sequence used here is based on sequence code kindly written and provided by Benedikt Poser. We thank Steve Cauley at MGH for sharing the interface of their image reconstruction for use with the SMS acquisition. We thank Miriam Heynckes for advice on the use of auditory stimulation setups. We thank Omer Faruk Gulban for early contributions to this work. We thank Chris Wiggins for providing the 3rd order shimming tools used here. Scanning was supported by FPN (Faculty of Psychology and Neuroscience) via the MBIC grant scheme. LKF was funded by the National Institute for Health grant RF1MH116978-01. FDM was funded by the National Institute for Health grant RF1MH116978-01 and the European Research Council (ERC) under the European Union’s Horizon 2020 research and innovation programme (grant agreement No. 101001270). Laurentius Huber was funded from the NWO VENI project 016.Veni.198.032.

## Data and Code availability

The anonymized raw data of this study is available and can be downloaded from https://doi.org/10.34894/WCUTUL. Fully processed data as shown in Fig. 2 are available for download here: https://layerfmri.page.link/A1VASO_3DMAPS. The scripts that were used in the analysis can be found on https://github.com/layerfMRI/repository.

## CRediT authorship contribution statement

**Lonike K. Faes:** Conceptualization; Investigation; Data curation; Formal analysis; Methodology; Project administration; Visualization; Writing - original draft. **Federico De Martino:** Conceptualization; Investigation; Funding acquisition; Supervision; Writing - original draft/review & editing. **Laurentius Huber:** Conceptualization; Investigation; Formal analysis; Methodology; Visualization; Writing - review & editing.

## Diversity Statement

Recent work in several fields of science has identified a bias in citation practices such that papers from women and other minorities are under-cited relative to the number of such papers in the field (Dworkin et al., 2020). In the human layer-fMRI community the average of the gender citation bias is 84% male, 15% female (https://layerfmri.com/papers/). We obtained the gender of the first author of each reference. By this measure (and excluding self-citations to all authors of our current paper), our references contain 78% male first and 22% female first. This method is limited in that: (i) names, pronouns, and social media profiles used to construct the databases may not, in every case, be indicative of gender identity, and (ii) it cannot account for intersex, non-binary, or transgender people. We look forward to future work that could help us to better understand how to support equitable practices in science.

