## Supplementary Material for "Cerebral blood volume sensitive layer-fMRI in the human auditory cortex at 7 Tesla: Challenges and capabilities"

**Supplementary materials**

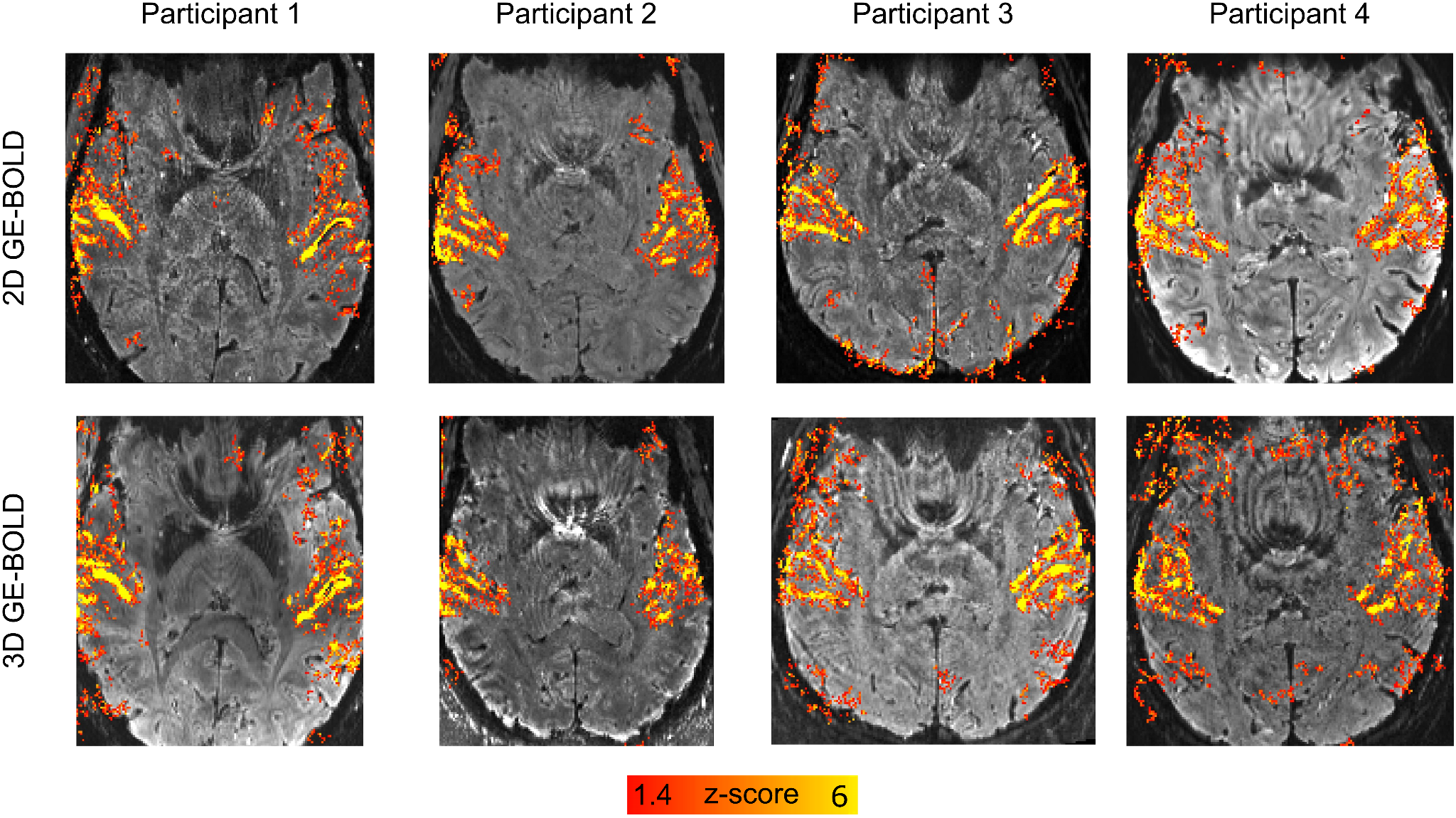

**Supplementary Figure 1.** Activation maps of BOLD (z-scores) overlayed on distortion corrected mean GE-BOLD EPI images (per participant and readout). The color map was chosen to match the VASO data displayed in figure 2.

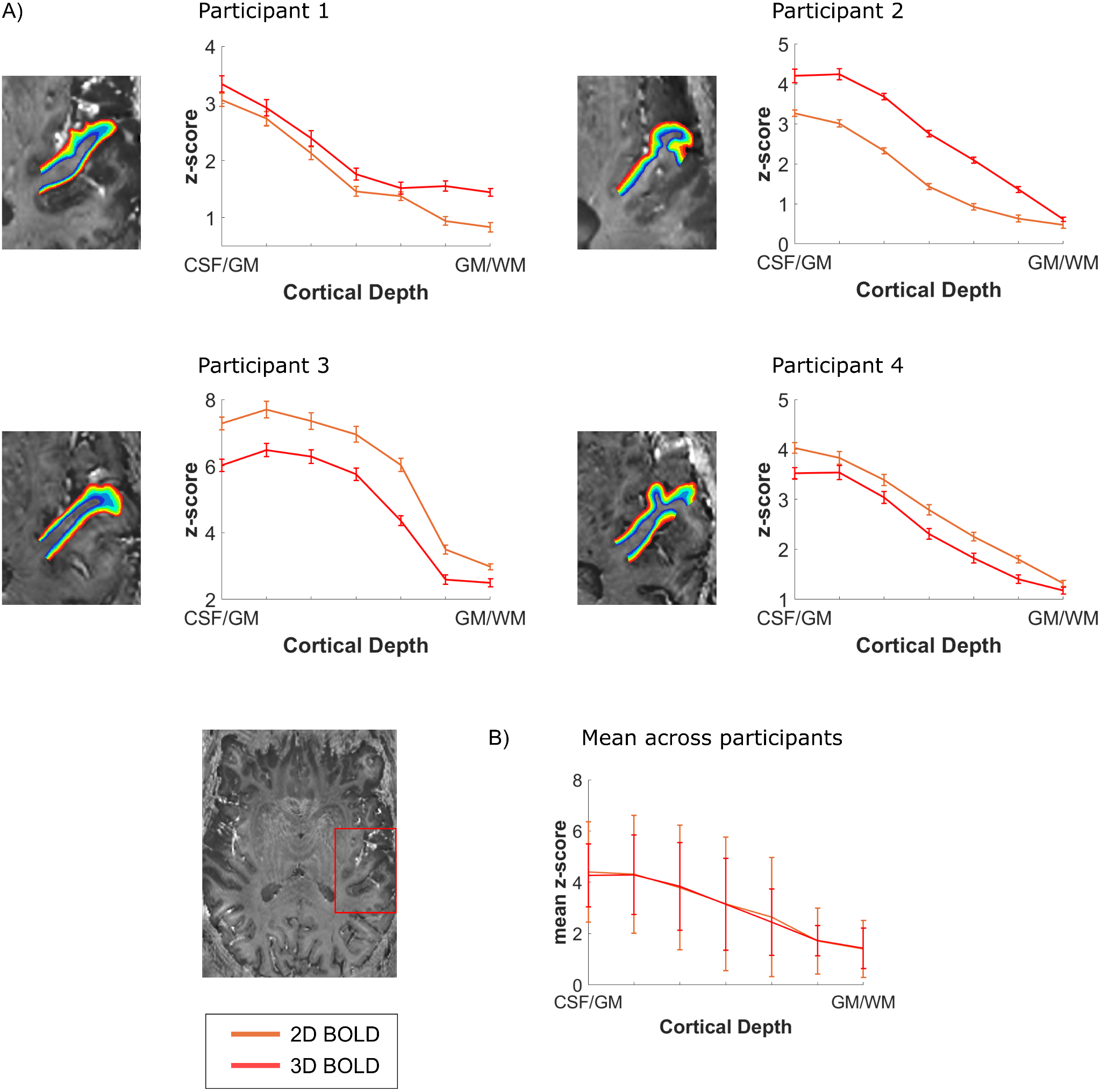

**Supplementary Figure 2.** Z-scored cortical depth-dependent activation changes for the 2D- and 3D-EPI BOLD data. A) Functional layer-dependent changes across depths for each participant. The BOLD data is coming from the same anatomically-based ROI that was used to calculate the layer-dependent VASO changes in figure 3. B) Average z-scored layer-dependent activation changes across participants.

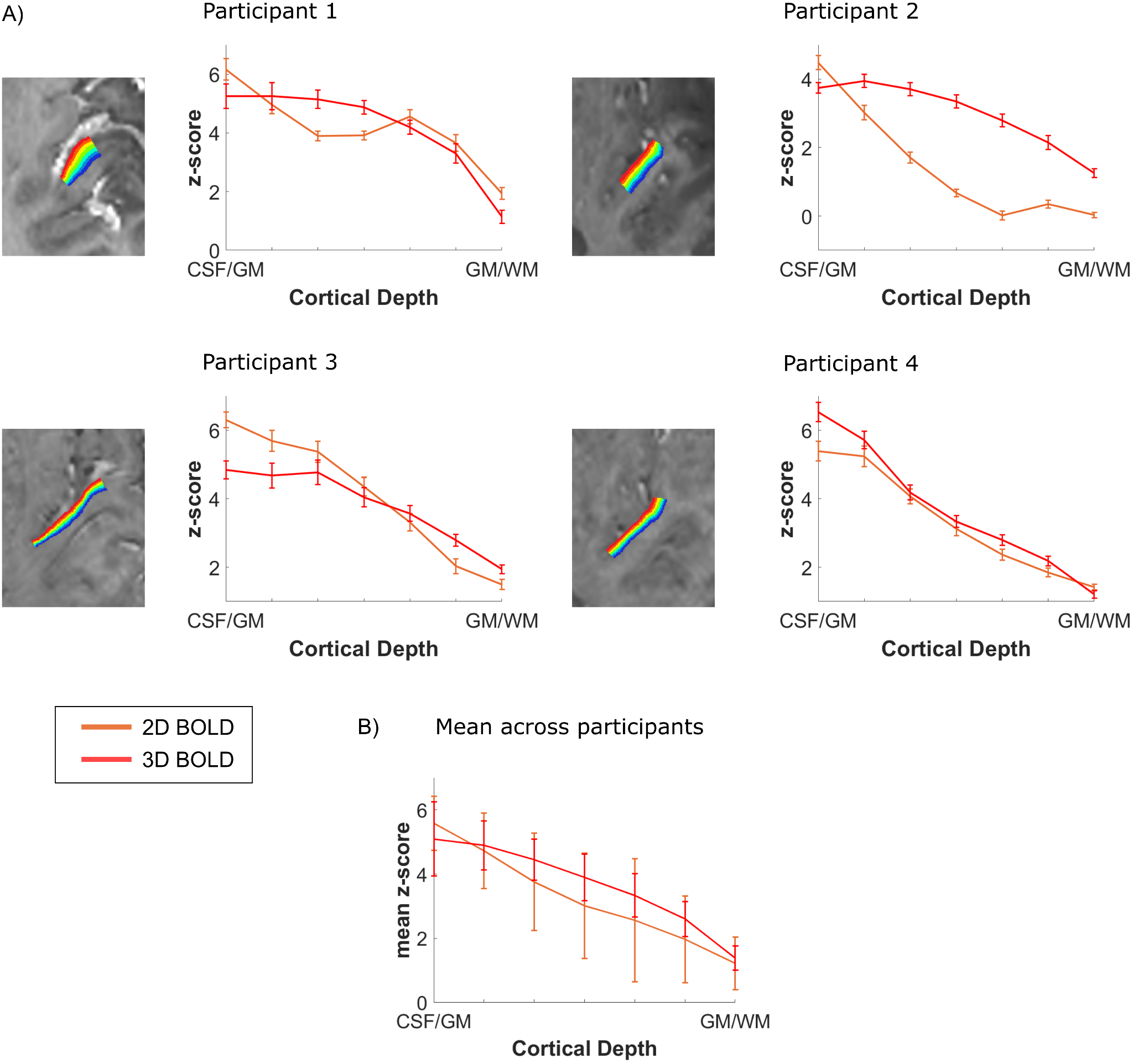

**Supplementary Figure 3.** Z-scored cortical depth-dependent activation changes for the 2D- and 3D-EPI BOLD data. A) Functional layer-dependent changes across depths for each participant. The BOLD data is coming from the same functional activation-based ROI that was used to calculate the layer-dependent VASO changes in figure 4, drawn on an axial slice as shown in figure 3A. A) B) Average z-scored layer-dependent activation changes across participants.

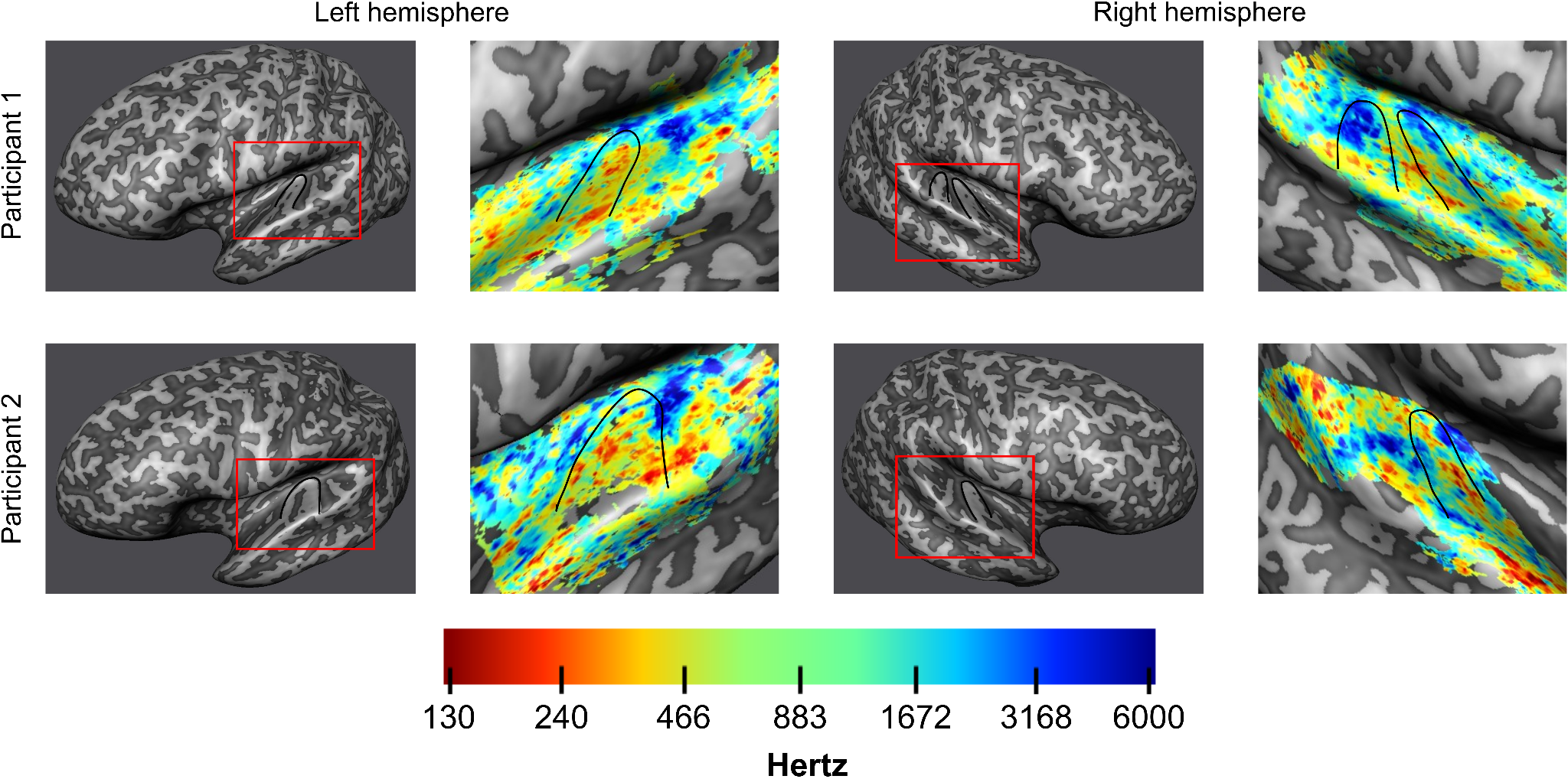

**Supplementary figure 4.** Inflated mid-gray matter surface meshes were created to visualize tonotopic maps created with the BOLD data. On the right of each inflated surface, tonotopic maps are displayed for both hemispheres of the two participants. Heschl’s gyrus is outlined in black. A tonotopic high-low-high frequency preference gradient is visible in the data.
